# Bimodal evolution of Src and Abl kinase substrate specificity revealed using mammalian cell extract as substrate pool

**DOI:** 10.1101/2020.08.12.248104

**Authors:** Patrick Finneran, Margaret Soucheray, Christopher Wilson, Renee Otten, Vanessa Buosi, Nevan J. Krogan, Danielle L. Swaney, Douglas L. Theobald, Dorothee Kern

## Abstract

The specificity of phosphorylation by protein kinases is essential to the integrity of biological signal transduction. While peptide sequence specificity for individual kinases has been examined previously, here we explore the evolutionary progression that has led to the modern substrate specificity of two non-receptor tyrosine kinases, Abl and Src. To efficiently determine the substrate specificity of modern and reconstructed ancestral kinases, we developed a method using mammalian cell lysate as the substrate pool, thereby representing the naturally occurring substrate proteins. We find that the oldest tyrosine kinase ancestor was a promiscuous enzyme that evolved through a more specific last common ancestor into a specific human Abl. In contrast, the parallel pathway to human Src involved a loss of substrate specificity, leading to general promiscuity. These results add a new facet to our understanding of the evolution of signaling pathways, with both subfunctionalization and neofunctionalization along the evolutionary trajectories.

## Introduction

The human genome contains 32 non-receptor tyrosine kinases (NRTKs) that are tightly involved in a multitude of cellular processes including differentiation, apoptosis, and proliferation^1-3^. The interaction between each NTRK and its substrate comprises a fundamental cellular signal that consequently required the evolution of specificity between signaling pathways. To prevent unwanted signaling ‘crosstalk’, NRTKs have evolved two main strategies to ensure substrate insulation^4,5^: kinase localization^6-12^ and active-site peptide specificity^13-18^. The localization process is achieved through binding interactions on the NTRKs’ SH2/SH3 domains, which complex with either phosphotyrosines or poly-prolines, respectively^19-22^. The differences in the active site of NRTK kinase domains results in specificity where only a subset of substrates can bind and thus get phosphorylated.

Unlike the serine/threonine family of kinases, NRTKs possess relatively promiscuous active-site peptide specificities with a broad range of potential substrates^5^. In high-throughput substrate screens, catalytic domains of NRTK members phosphorylated hundreds of distinct peptide sequences, highlighting the promiscuity of these kinases. Nevertheless, comparisons within the family show unique sequence preferences and a consequent range of substrate selectivity. For two of the most well-studied members, Abl and Src, narrow and broad selectivities are reported, respectively^13,17^. Because substrates bind at the active site in an elongated fashion, the primary peptide sequence largely dictates the description of selectivity^5^. Abl has a clear preference for hydrophobic residues flanking the tyrosine of interest (I/L/V_-1_, A_+1_, and P_+3_)^14,23^. In contrast, little sequence selectivity is observed for Src other than the relatively weaker preferences for a bulky aliphatic residue (I/V/L_-1_) at the residue preceding the phosphoacceptor, a phenylalanine three residues away from the phosphoacceptor (F_+3_), and a negatively charged residue on the N-terminal side of the phosphoacceptor (D/E_-4, - 3, −2_) ^17,24,25^.

As Src and Abl are sister clades within the NRTK phylogeny, their distinct sequence preferences beget the question: how did peptide selectivity arise throughout evolution? Herein we answer this question using ancestral sequence reconstruction (ASR) for the catalytic domains. ASR uses modern sequences and an evolutionary model to infer the sequences of internal nodes in a phylogenic tree (Figure 1A, Figure 1 — figure supplement 1) ^26-31^. Resurrection of ancestral kinases bridging Src and Abl allows the evolution of peptide selectivity to be directly traced. The differences in protein sequence between modern kinases and ancestors are spread through the kinase domain and the oldest ancestor only has ∼65% similarity with Src (Figure 1B).

**Figure 1.**
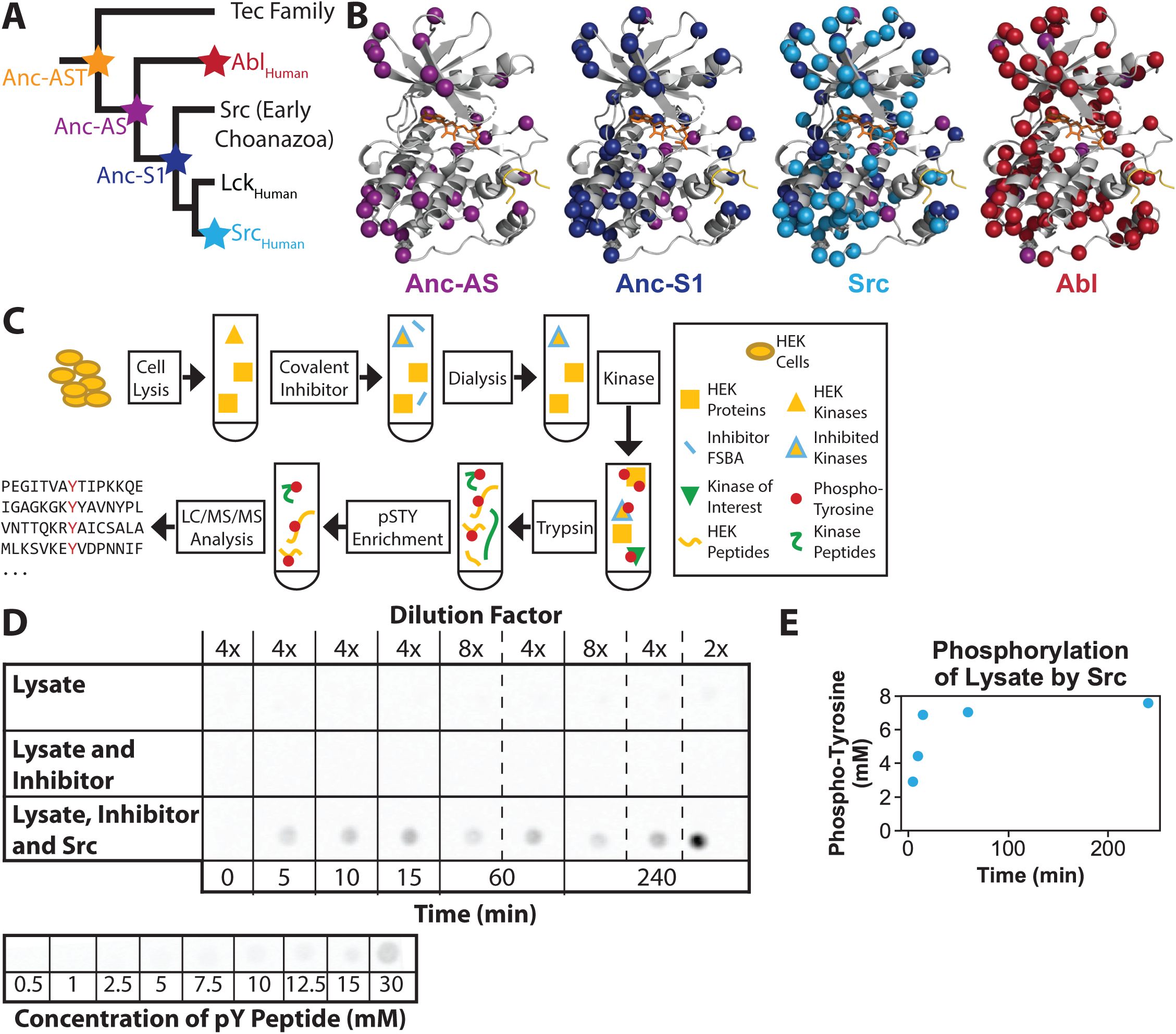
Analysis of the evolution of non-receptor tyrosine kinase specificity with HEK lysate based approach. **(A)** Phylogenetic tree of non-receptor tyrosine kinase domains constructed with Bali-Phy (Suchard *et al*. 2006). The reconstructed nodes and modern kinases used herein are marked with stars. For complete tree, sequences, testing of alternate sequences, and statistics see (Wilson *et al*. 2015). **(B)** Crystal structure of Abl (PDB ID: 2G2I; Levinson *et al*. 2006) with peptide substrate (yellow) and ADP (orange) bound. The additive differences in primary sequence between Anc-AST and Anc-AS (purple; 87.3% identity), Anc-S1 (dark blue; 85.8% identity), Src (blue; 65.2% identity), and Abl (red; 67.0% identity) are shown. **(C)** Flow chart displaying how cell lysate was prepared and used for kinase specificity assays. See methods for a detailed description. **(D)** Western dot blots using anti-pY1000 antibody to detect phosphorylated tyrosine in proteins. (Top) Cell lysate incubated without Src does not show any phosphorylation, whereas lysate treated with the kinase does. Dilutions of the 60-minute and 240-minute time points illustrate that measurements are in the linear range of the dot-blot. (Bottom) Control of phosphorylated peptide blotted at a range of concentrations, diluted 4-fold. **(E)** Phosphorylation of the cell lysate over time by Src kinase.

Src and Abl display lower sequence selectivity than other kinases such as Aurora A and B-RAF serine/threonine kinases^5,24,32^, and, consequently, obtaining an accurate description of the primary sequence determinants for each tyrosine kinase is a greater statistical challenge^33^. Comparison of ancestral and modern kinases requires a comprehensive library of substrates since ancestral kinases likely refined their sequence preferences over time. To ensure biological relevance, the peptide library should ideally be composed of naturally occurring proteins. To construct such a library, we took advantage of the diversity of sequences present in mammalian whole cell lysate (HEK293 cell line)^34,35^. After endogenous kinases are covalently inhibited, the proteome of mammalian cells presents a convenient substrate library containing thousands of potential protein substrates. Here we use this comprehensive library to examine the evolution of substrate selectivity in Abl/Src tyrosine kinases. We find that kinase substrate preferences evolved in a complex manner involving two different modes: a promiscuous progenitor specialized into the modern specific Abl, whereas evolution of Src involved relaxing selectivity via a specific ancestral intermediate. We find that kinase substrate preferences evolved in a complex manner involving two different modes: a promiscuous progenitor specialized into the modern specific Abl (subfunctionalization), whereas evolution of Src involved relaxing selectivity via a specific ancestral intermediate (neofunctionalization). Therefore, our results shed light into a critical open question in signaling, how new protein kinases with novel substrate specificities have evolved.

## Results

### Whole-cell lysate phosphoproteomics-based approach

A large and diverse protein library is necessary to readily screen the primary sequence determinants of ancestral and modern NRTKs. An easily accessible, cheap, and biologically relevant pool of substrates was created by inactivating endogenous kinases in HEK293 lysate using the covalent, nonspecific inhibitor 5′-[p- (fluorosulfonyl)benzoyl]adenosine (FSBA) (Figure 1C). After dialyzing out unreacted inhibitor, purified kinases were added to the treated lysate and phosphorylation was initiated with the addition of Mg^2+^ and ATP. To assess the required reaction time, total phosphorylation was monitored by western dot blot experiments. Constant phosphorylation levels were found to occur between two to four hours (Figure 1D,E). After protein digestion with trypsin, peptide fragments were enriched for phosphorylation using immobilized metal affinity chromatography (IMAC) and then analyzed by liquid chromatography-mass spectrometry (LC/MS/MS, Figure 1C). Peptides were associated with their full protein sequence based on the known HEK293 proteome, and results were focused on a 15-amino acid sequence window centered on the phosphorylated tyrosine.

To determine a kinase’s sequence specificity, each amino acid frequency must be compared between the phosphorylated dataset and the background HEK293 proteome^33^. A background dataset was generated by proteomic analysis of lysate that was treated with kinase inhibitor, but where no kinase was added and no phosphorylation enrichment was performed^36^. From this analysis, the amino acid frequencies of all surrounding tyrosine residues (not only phosphorylated tyrosines) were calculated (Figure 1 — figure supplements 2 and 3).

To determine the extent of endogenously phosphorylated peptides present, we analyzed a lysate sample that was enriched for phosphorylated peptides but lacked exogenous kinase. A total of 26 phosphorylated tyrosines were found in this control and these peptides were excluded from the ancestral/modern NRTK list of substrates in which kinase was added (Figure 1 — figure supplement 4).

In the data set where Src was added to the cell lysate, 8208 unique phosphorylated sequences were identified (Figure 2A). These peptides include the characteristic preferences for large aliphatic residues directly preceding the phosphotyrosine (V_-1_), negatively charged residues in multiple positions N-terminal to the phosphotyrosine (D/E_- 3, −2_), and a glycine following the phosphorylation site (G_+1_). An inclination for other aliphatic residues preceding the phosphotyrosine (I/P/T_-1_) is also seen (Figure 2B). Notably, a preference for proline at the −1 position identified in our data was not observed previously. For Abl, specificity for large aliphatic residues preceding the phosphotyrosine is found (I/V_-1_), as well as the canonical proline at the +3 position (Figure 2B). Additionally, Abl exhibits a high preference for proline at the −2 position, which had not been identified previously.

**Figure 2.**
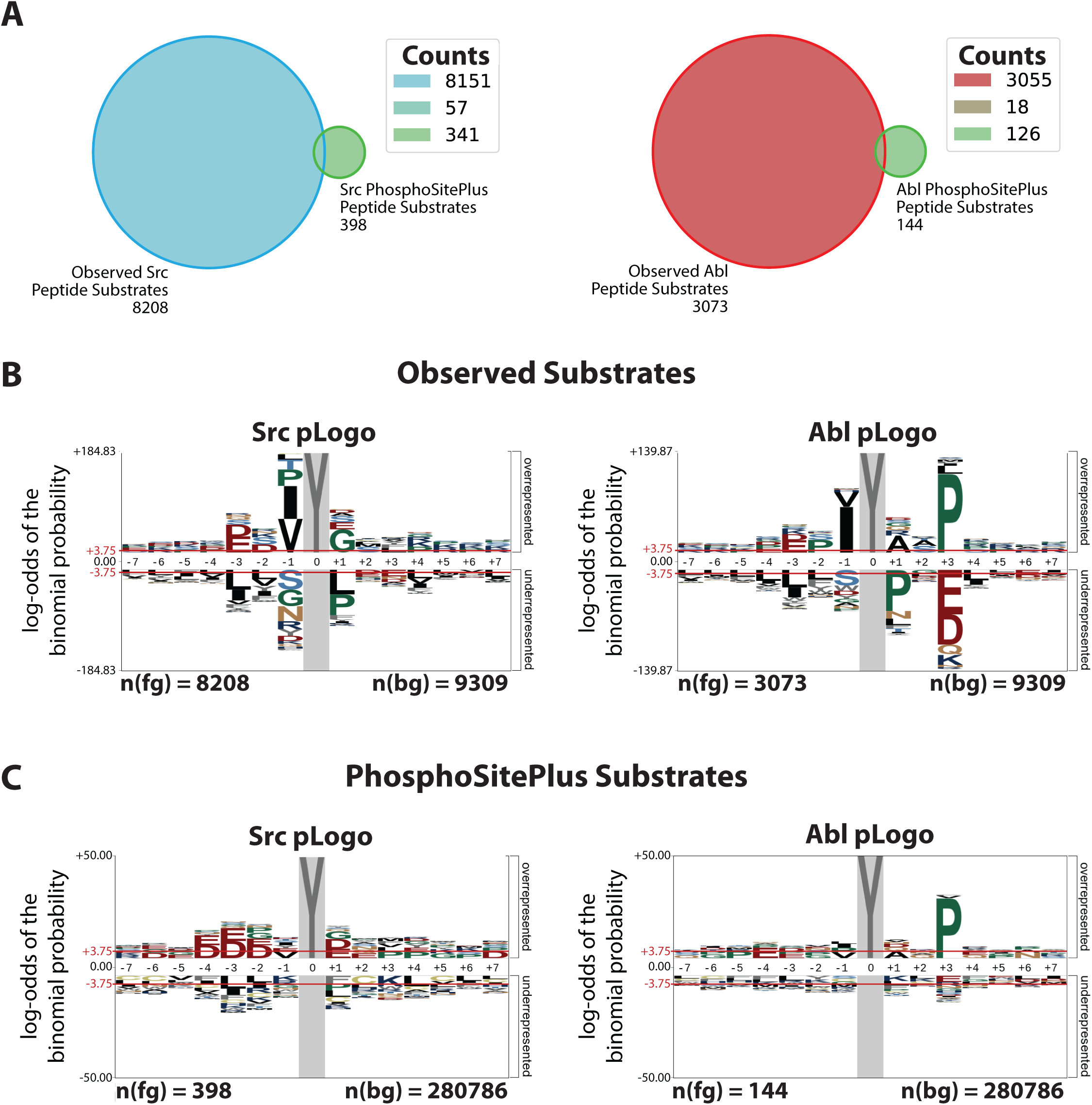
HEK cell lysate library approach finds large number target substrates and recapitulates known specificities of Src and Abl with improved statistics. **(A)** Venn diagrams of sequences that were observed in our Src and Abl kinase specificity experiments compared to the substrates listed in PhosphoSitePlus (Hornbeck *et al*. 2015) for the respective kinase. **(B)** pLogo’s for the prevalence of each amino acid in Src and Abl’s individual set of substrates (8208 and 3073 sequences, respectively) relative to the background experiment (9309 sequences). **(C)** In comparison, pLogo’s generated from the list of human substrates on the PhosphoSitePlus database (Hornbeck *et al*. 2015). Statistical significance level is shown as red line in B and C.

We can compare the results obtained here with the substrate specificity that is observed when only natural substrates are considered. PhosphoSitePlus is a database which annotates all known phosphorylation sites *in vivo* and *in vitro* that a given kinase phosphorylates^24^. Overall, our results confirm the previously found descriptions for both Src and Abl’s substrate specificities based on substrates in the PhosphoSitePlus database, but additionally identify a few new preferences (Figure 2B, C). Our HEK293 lysate has a much larger number of phosphorylated substrates than the PhosphoSitePlus database, which allows us to ascertain the residues dictating phosphorylation specificity with greater accuracy and statistical significance than is possible with the PhosphoSitePlus data (Figure 2B,C). Some of the differences may be due to the larger number of substrates in our whole cell lysate. However, many of the observed discrepancies likely result from differences in experimental design. PhosphoSitePlus is based on *in vivo* substrates for the full-length kinase, whereas we are interested in the intrinsic specificity of the kinase domain. In our experiments, we do not have the full-length kinases and, therefore, we only find substrates that are selected by the kinase domain itself. In contrast, phosphorylation within the cellular framework, as reported by PhosphoSitePlus, is strongly determined by regulation and co-localization events, and intrinsic kinase domain specificity plays a relatively smaller role. Indeed, Shah *et al*. studied the specificity of NRTK kinase domains with a high-throughput, cell-surface based experiment and found similar discrepancies between PhosphoSitePlus based logos and their experimentally determined sequence determinants^17^.

### Evolution of specificity between Src and Abl

Having established now the accuracy and statistics of our methodology on the modern kinases, we next chose to determine the sequence specificity of three resurrected ancestral kinases (Figure 1A). Anc-AS and Anc-S1 were previously resurrected for investigating the mechanism of Abl selectivity for Gleevec^29^, while the newly resurrected Anc-AST (Figure 1 — figure supplement 1) is the oldest common ancestor of the Abl/Src branch and the Tec family. Using our whole cell lysate phosphoproteomics-based method, we identified a total of 12,056 unique sequences phosphorylated by the ancestral and modern proteins (Figure 3A, Figure 3 – figure supplement 2). The common ancestor of Src and Abl, Anc-AS, phosphorylated the least number of substrates (2495), which was comparable to Abl (3073). The relative dearth of substrates for Anc-AS hinted that this ancestor might be more specific than the promiscuous Src, which phosphorylated a total of 8208 substrates. In contrast, the ancestors preceding (Anc-AST) and following (Anc-S1) the common ancestor of Src and Abl each phosphorylated a significantly greater number of substrates (8189 and 7242, respectively), indicating these ancestors were likely more promiscuous.

**Figure 3.**
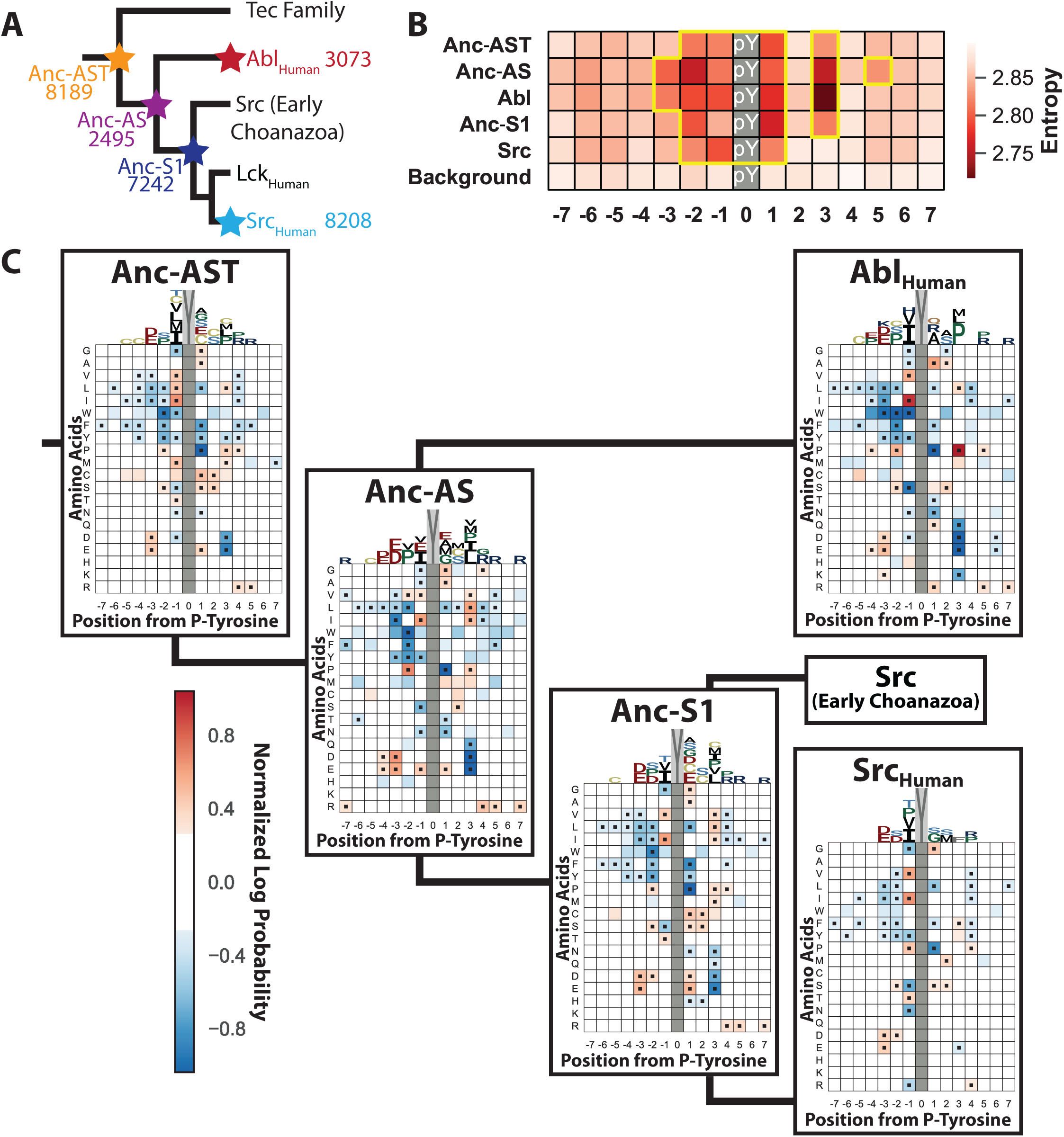
Anc-AST and Anc-S1 are promiscuous, but are bridged by a relatively specific Anc-AS. **(A)** Total number of phosphopeptides found for each of the five kinases are plotted onto the gene tree. **(B)** Positional entropy description for each kinase, where a lower entropy indicates higher specificity. Highlighted in yellow are values below the ∼30th percentile for sequence entropy, illustrating the positions with the highest specificity. **(C)** 20×15 positional/amino acid heatmap displaying the normalized log probability of each amino acid at a given position for modern and ancestral kinases. Significant residues at each position are marked with black squares (p<0.05). The sequence logo above the heatmap displays positions with positive values only, and the height of the character is equal to the normalized log probability.

General kinase specificity was then quantified by calculating the substrate sequence entropy for each position in the 15-residue window (Figure 3B)^37^. Lower entropy, with fewer potential amino acid possibilities, indicates higher specificity. As expected from the sequence logos (Figure 2B), Abl possesses the lowest entropy at residues close to the phosphorylated tyrosine (−3, −2, −1, +1, and +3), with the most specificity occurring at the signature proline position (+3). In contrast, the promiscuity of Src manifests itself as increased entropy across almost all positions and a more limited region of high specificity (−2, −1, and +1). The ancestral proteins give entropy plots that agree with what was suggested by the observed substrate counts: The common ancestor (Anc-AS) possessed entropy akin to Abl, albeit with a higher entropy at +3 and lower entropy at positions −2 and +5. The two additional ancestors, Anc-AST and Anc-S1, both exhibited ‘hybrid’ specificity with higher entropy than Abl, but less than Src. Notably, only Src lacks specificity at the +3 position.

To analyze each enzyme’s positional specificity in more detail, specificity heat maps were created to illustrate the relative specificity for each amino acid at every position in the 15-residue window (shown as a 20×15 matrix of positional normalized amino acid log probabilities, Figure 3C)^17,38^. Qualitatively, Abl’s specificity is apparent from the high-intensity signals for both preferred residues (red, P_+3_ and I_-1_) and unfavorable residues (blue, S_-1_, P_+1_, and D/E_+3_), while Src has more white space and overall less intense signal. We note that under substrate saturating conditions the enzyme would phosphorylate even the less favorable substrates, which could result in an apparent low specificity. To ensure that the observed promiscuity is not due to substrate saturation, experiments with Src were repeated with a much shorter incubation period of the cell extract and the kinase (10 minutes versus 4 hours). In this control, less phosphorylation was observed (Figure 1 C,D); however, the same primary sequence determinants were found (Figure 3 — figure supplements 1 and 2), validating our findings.

Tracing individual amino acid preferences at specific positions provides a clear picture of how specificity evolved for Abl (Figure 4A). Focusing on residues which are preferred in Abl, but disfavored in Src (P_+3_, A/V_+2_, A_+1_, and P_-2_), we see that moderate preference is already observed in ancestral kinases. The evolutionary path from the oldest ancestor (Anc-AST) to Abl involves further increasing specificity either at the Anc-AS node (P_-2_ and A/V_-1_) or in the final transition to Abl (A_+1_ and P_+3_). In contrast, the pathway from the more specific Anc-AS to Src involves a corresponding loss of specificity for each of these residues, with Anc-S1 possessing intermediary preferences (Figure 4A).

**Figure 4.**
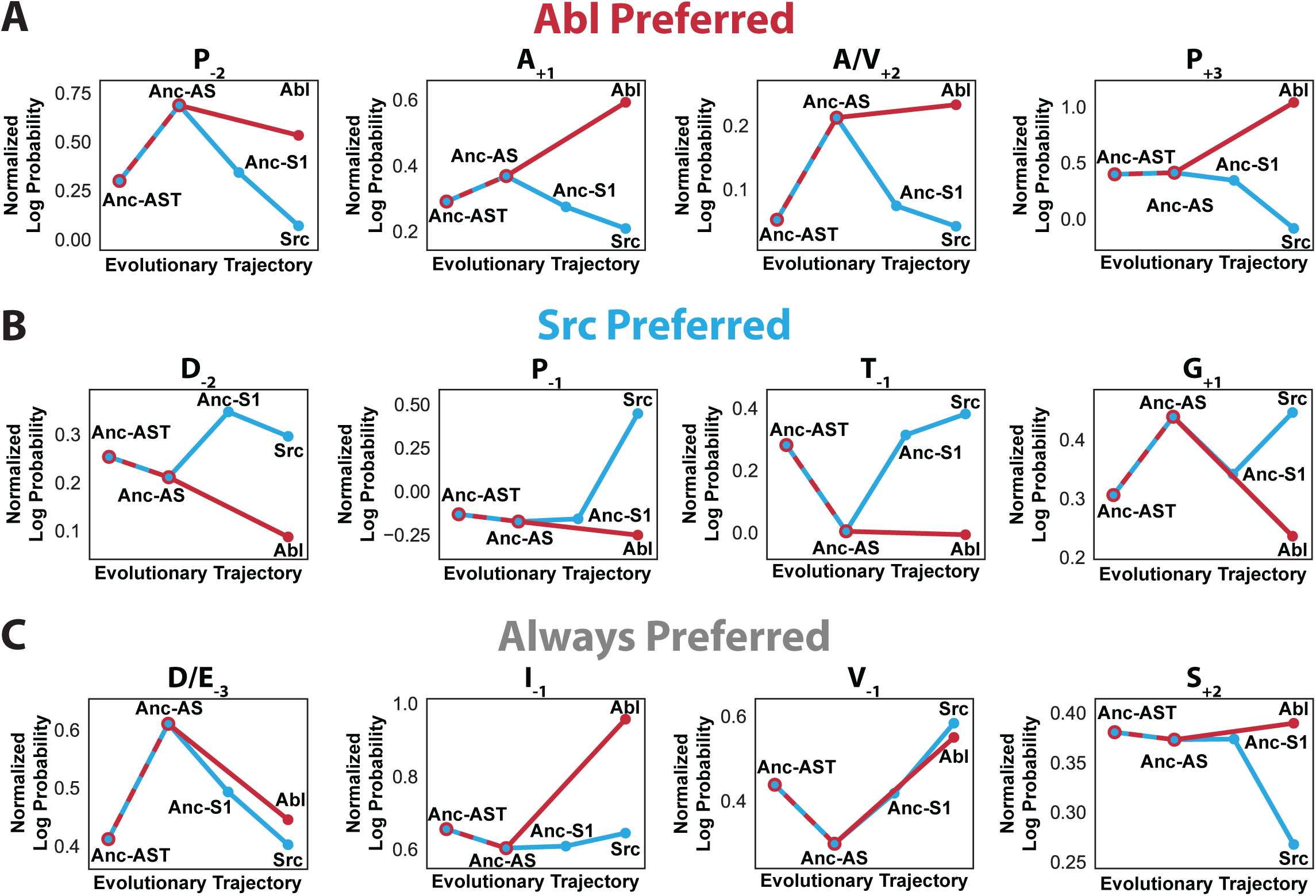
Evolutionary trajectories of sequence specificity show both subfunctionalization and neofunctionalization. The normalized log probability of an amino acid occurring in a pool of substrates demonstrates the evolutionary progression for **(A)** Abl specific residues, **(B)** Src specific residues, and **(C)** residue determinantscommon to all all kinases in this study.

The evolution of the few positions favored by Src followed a different trend. The preference for P_-1_ appears late, only in Src. Such recent evolution is in agreement with the lack of preference for proline at position −1 of its close homolog Lck^17^. Other Src sequence preferences were already present in the oldest ancestor, then lost in Anc-AS and regained in Anc-S1 and Src (Figure 4B).

The increased promiscuity of Src is revealed in overall lower log probability values than that of Abl’s specific residues. Despite these differences in substrate specificity, there are multiple positions where all modern and ancestral kinases prefer the same residues, most of which are well-known features of NRTKs (e.g., I_-1_, D/E_-3_, and S_+2_) (Figure 4C). As these characteristics are observed in all ancestral and modern proteins in our study and are common among most NRTKs, they likely represent the oldest features of substrate specificity for the NRTK family.

### Validation of evolutionary trends of primary sequence determinants via enzyme kinetics

Having determined the primary sequence determinants for the ancestral and modern kinases, *in vitro* peptide enzyme turnover experiments were performed to relate these bulk specificity experiments to quantitative enzymatic parameters. Compressing thousands of substrates into residue-by-residue descriptions is compelling (i.e., preference for P_+3_), but how these preferences relate to enzymatic properties remains unclear. We therefore measured the Michaelis-Menten kinetics of four distinct peptide substrates with each of the five ancestral and modern kinases.

Previous microarray specificity experiments had determined optimized substrates for Src and Abl, known as Srctide and Abltide, respectively^13,18,25^. While both substrates are ideal for their respective modern kinases, each also has residues favored by the opposite kinase, which allow it to be phosphorylated to a certain degree by each kinase. Therefore, we designed modified versions of Srctide and Abltide, called Srctide2 and Abltide2, to test our evolutionary trends (Figure 5A,B). Srctide2 was intended to be favored by both Src and Anc-S1, with changes made to Srctide to include residues that occurred more frequently in substrates for these two kinases (D_-2_, A_+2_, and I_+3_). Abltide2 was designed to be preferred by both Abl and Anc-AS by mutating the alanine at position −2 into a proline. P_-2_ is favored by all the kinases, except for Src (Figure 5A, B).

**Figure 5.**
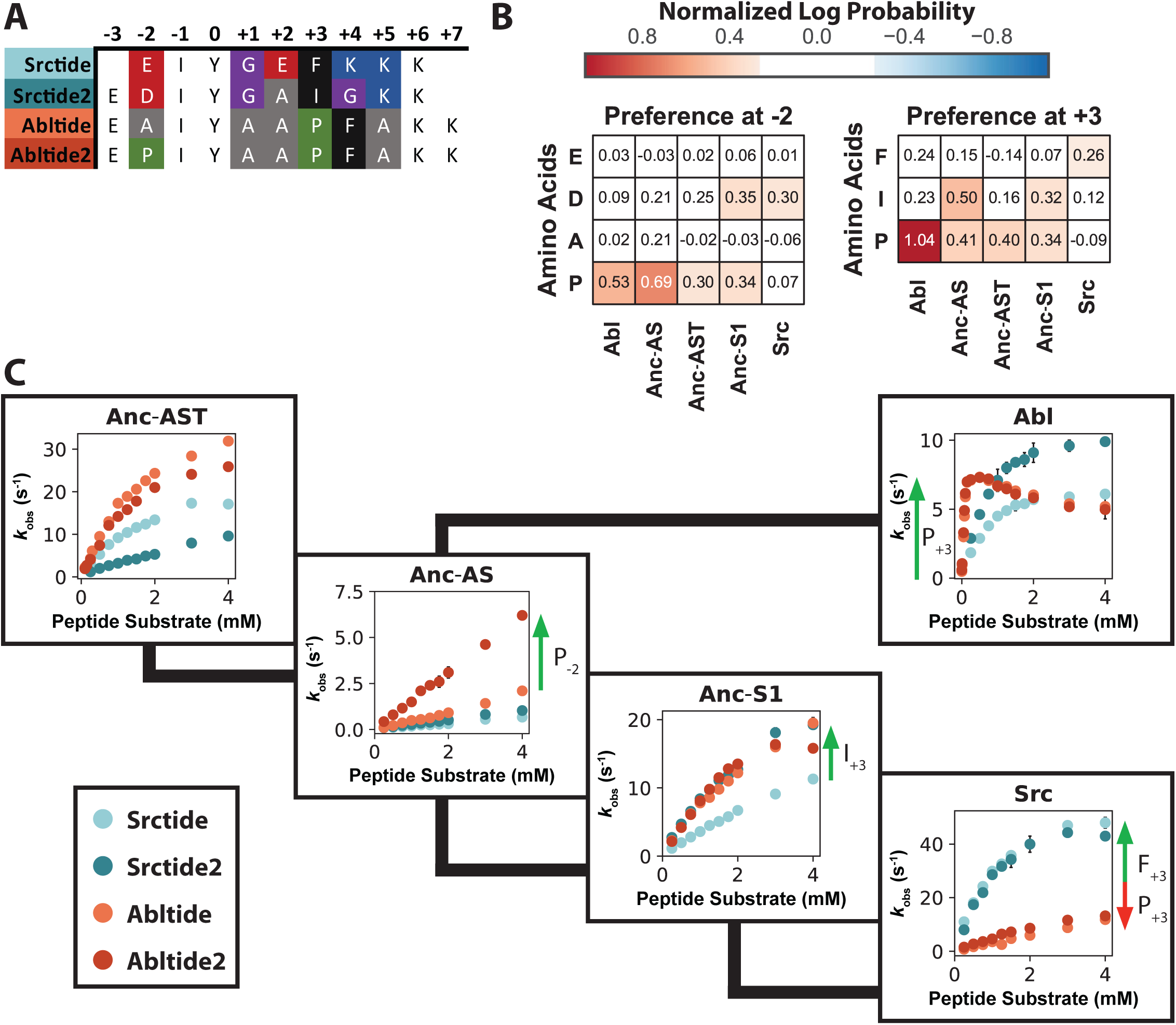
Specificity of kinases correlate with individual peptide turnover parameters. **(A)** Sequence alignment of the substrate peptides used with differences between peptides highlighted. Coloring of the peptides are used in the Figure 5C. **(B)** Normalized log probability of amino acid preferences for −2 and +3 positions for the five ancestral and modern kinases determined from the cell lysate experiments, see also Fig. 3C. **(C)** Michaelis-Menten curves of phosphorylation for the four peptide substrates and the five different kinases measured with a coupled assay for detecting ADP production (see methods). While it is not possible to saturate and measure accurate *K*_*M*_ values for many of the substrates, it is possible to observe how the rates are affected under *k*_*cat*_*/K*_*M*_ conditions. Each kinase was assayed at 20-200 nM, and error bars represent the standard deviation from three measurements. Green arrows indicate the key primary sequence determinants in the different substrates responsible for the large changes in observed rates.

As substrates in signaling cascades are generally present at low concentrations *in vivo*, the *k*_cat_*/K*_M_ likely represents a more fundamentally important parameter than *k*_cat_ for substrate specificity. As can be seen from the measured Michaelis-Menten curves (Figure 5C), the measured differences in the kinetics corroborate the evolutionary trends found before but suggest additional features for substrate specificities. Starting with a promiscuous Anc-AST, Anc-AS becomes more selective, particularly for substrates with P_-2_, Ablitide2 (which was identified as a highly preferred residue for Anc-AS and Abl from our phosphoproteomics data, Figure 5B). Moving to Abl, the specificity for Abltide and Abltide2 further increases as seen with high increases in *k*_cat_*/K*_M_. This is primarily due to the strong preference for P_+3_ (Figure 5B), which are both present in Abltide and Abltide2. At higher concentrations of these two well-optimized peptides we observe partial inhibition, due to the negative cooperativity of ATP and peptide substrate found for Abl^39^. Following the evolutionary branch towards Src, Anc-S1 becomes more promiscuous, mainly due to its ability to catalyze Src-preferred substrates in addition to the Abltide substrates. Furthermore, the strong preference of I_+3_ observed in the proteomics data (Figure 5B) can be directly recapitulated by the preference for Srctide2 (Figure 5C). The strong preference of Srctide and Srctide2 over the Abltide substrates only appears in Src, primarily due to a complete loss of preference for P_+3_ leading to poor activity for the Abltide substrates, combined with a subtle preference for F_+3_.

## Discussion

There are multiple methods described in the literature for assessing substrate specificity of protein kinases, including non-receptor tyrosine kinases and serine/threonine kinases. Several studies look only at known natural substrates, such as in PhosphoSitePlus, to determine which residues occur more frequently in known phosphorylation sites^24,40^. Other studies, including the one presented here, use the whole cell lysate as a pool of substrates to test the specificity of kinases^34,35^. The HAKA-MS method reported by Muller *et al*. used a similar whole cell lysate, yet only found a P_+3_ preference from 104 Abl substrates (Figure 2 – figure supplement 2)^35^. In contrast, our study found ∼30-fold more substrates and several residues that are preferred or disfavored at multiple positions. The large difference in detected phosphorylated substrates could be due to different methods to enrich for phosphorylated peptides: Muller *et al*. use multiple phosphotyrosine binding antibodies, whereas we use IMAC. We also tried using phosphotyrosine binding antibodies for enrichment, but we found peptide bias and artifacts using this method. Lastly, one of the most popular methods uses peptide libraries to determine the preference of kinases^13,14,16-18,38^.

Our experimental method, which exploits the substrate-rich, kinase-inactive whole cell lysate and has improved statistical significance, discovered new determinants of Abl and Src specificity in addition to those previously reported^17^: Src shows significant favorability for proline or threonine at the −1 position and a serine at the −2 location, while Abl shows a preference for proline at the −2 position and a serine at the −2 position. These new features were subsequently verified by *in vitro* enzyme kinetic experiments. The most important advantage of the increased sensitivity of our assay has been the ability to explore how Src and Abl kinases evolved their sequence preferences.

How these kinases have differing specificities can be partially rationalized based on the details of kinase sequence and structure. Y569 in Abl has previously been determined to be required for its preference for proline in the +3 position of substrates^41^. Leucine at the homologous position (L475) in Src was previously shown to disfavor proline^41^. A437 has been proposed to be responsible for Src’s preference for phenylalanine at position +3, along with L475^17^. All ancestors contain an isoleucine at position 475 and show intermediate specificity towards P_+3_. Moreover, Anc-S1 differs from Src in both positions, I475 and L437 (Figure 5 – figure supplement 1), identical to another Src family kinase member, Lck (I412 and I450 in Lck). Interestingly, when comparing the Anc-S1 substrate preference to that of Lck, as investigated by Shah *et al*.^17^, we see a high similarity in the +3 position preferences. Both Anc-S1 and Lck show a strong preference for L_+3_ and P_+3_, suggesting that Anc-S1 is more Lck-like, and that the substitution to the less bulky A437 in Src causes its preference for F_+3_ (Figure 5 – figure supplement 1). These structural differences between Src and the ancestors explain why the Srctide is less effective at being phosphorylated by other kinases. Anc-S1 was unable to effectively phosphorylate Srctide potentially due to F_+3_, wherein its substitution to the less bulky isoleucine in Srctide2 is favored by Anc-S1. We elect to steer clear from additional structural explanations for other detected specificities, as these are just coarse models of kinase/substrate complexes, and more collective, long-range effects often underlie such specificity changes. For example, in an appealing study of the evolution of CMGC kinases, Howard *et al*. identified a key residue for imparting specificity at the +1 position^28^. Tests of their hypothesis via mutations in the corresponding modern kinases resulted in partial changes in specificities. Since the authors were unable to achieve a full swap in specificity, they concluded that there must be additional residues in play that are not readily apparent by looking at the differences in active-site residues.

The different trajectories we find in the evolution of Src and Abl substrate specificity add a new facet to our understanding of the evolution of signaling pathways. It has been postulated that much of biological diversity, including metazoan complexity, has been driven by the evolution of new regulatory networks and signaling pathways, such as those controlled by the post-translational modification of kinase phosphorylation^42,43^. A key evolutionary challenge in creating a new signaling pathway is ensuring kinase specificity to minimize crosstalk with other pathways, many of which are vital to cellular fitness. One critical open question is how new protein kinases with novel substrate specificities have evolved.

Gene duplication has been the major force driving the evolutionary diversity of signaling pathways and kinase specificity^42-44^. There are two main ways that gene duplication can evolve enzymes with new and different functions: (1) “subfunctionalization”, the specialization of previously existing functions, and (2) “neofunctionalization”, the creation of a novel function through the accumulation of beneficial, gain-of-function mutations^45,46^. The evolution of sequence specificity does not cleanly fall in either of these categories, because specificity is not an “all-or-nothing” gain or loss of a function. Nevertheless, for kinases, subfunctionalization most closely aligns with evolution from a non-specific ancestor (which can bind and use many different substrates) to descendants with differing, specialized specificities (which can bind and use only a subset of the ancestral substrates), whereas neofunctionalization involves evolution from a specific ancestor to a descendant with a broad, promiscuous specificity or with a different specificity. Though neofunctionalization is the older of the two hypotheses^47,48^, it is now widely viewed as relatively improbable and hence less frequent in evolution than subfunctionalization mechanisms^49-53^. For instance, in the evolution of specificity in CMGC protein kinases, the ancestral kinase was a promiscuous bispecific enzyme in respect to the +1 position, unlike the modern kinases which are specific for a single amino acid^28^. Currently, the evolutionary mechanisms by which gene duplications evolve new functions are controversial, and there are relatively few examples of classic neofunctionalization ^54-56^.

Intriguingly, we see both mechanisms of gene duplication in the evolution of Abl and Src. With Abl, subfunctionalization converted a promiscuous, non-specific ancestor (ANC-AST) into specific descendants (ANC-AS and modern Abl); with Src, neofunctionalization transformed a surprisingly specific ancestor (ANC-AS, the last common ancestor of the Abl and Src kinase families) into progressively more promiscuous descendants (ANC-S1 followed by modern Src). In fact, the particular lineage leading from ANC-AST to modern Src appears to involve both mechanisms.

In the toxin-antitoxin signaling systems of bacteria, Aakre *et al*.^57^ found that the evolution of some enzymes passes through promiscuous intermediates before developing strong substrate specificity. Our results suggest that eukaryotic protein kinases similarly evolve through waves of promiscuous and specific effectors. Periods of increased promiscuity may allow kinases to access new substrates while maintaining their current function, which could explain the redundancy of kinase networks^24^. To insulate individual pathways from others, sometimes this promiscuity may be selected out. In other cases, the overlap may provide a fitness advantage resulting in modern protein kinases with overlapping substrates.

The present work has addressed one component of kinase substrate specificity – the intrinsic specificity of the kinase domain – but it is important to emphasize the rich literature about the crucial role of the additional regulatory domains found within most NRTKs for kinase specificity. Proteins containing SH2 or SH3 domain binding sites are more likely to be phosphorylation targets for NRTKs, due to the selective activation of the kinases^7,20,58-60^. SH2 and SH3 domains have their own unique specificity^61-65^, and bring the kinases to their target substrates. Studies using kinase domains in isolation have not identified all the known substrates that are found in databases like PhosphoSitePlus. For example, Shah *et al*. limited their library to only known phosphorylation sites, yet they could not see a preference for all known kinase substrates for a given kinase^17^. We fully agree with their discussion^17^ that intrinsic kinase domain specificity^13,14,25^ acts in concert with the selective activation and localization provided by the SH2 and SH3 domains in a cellular context ^4,7,8,12,20,21,64,66-70^ to provide the full specificity of the NRTKs.

## Supporting information

Figure 3 - figure supplement 2: PhosphoproteomicsData.csv

Figure 3 - figure supplement 3: PhosphoproteomicsSrcShortTimePoint.csv

Figure 1 - figure supplement 3: BackgroundUnenrichedProteomics.csv

Figure 1 - figure supplement 4: ControlPhosphoproteomicsData.csv

## Materials and Methods

### Kinase Specificity in Whole Cell Lysate Experiment

HEK293 cells were grown in CytoOne 150×20mm TC dishes with DMEM (High Glucose, No Glutamine; Fisher Sci) containing HyClone bovine growth serum (Fisher Sci), glutamine (Fisher Sci), fungizone (Fisher Sci), and penicillin-streptomycin (Fisher Sci). At ∼90% confluency, cells were washed with 5mL of PBS before being harvested. Cells were centrifuged at 4500g for 6 minutes to pellet. PBS was then decanted from the pellet. Pelleted cells were washed by repeating the previous step. The pellet was then resuspended in ∼3 mL of assay buffer (20 mM Tris, Fisher; 500 mM NaCl, Fisher; 1 mM MgCl_2_, Fisher; 1mM TCEP, Fisher; pH 8) per 10 plates harvested. Cells were lysed by sonication followed by centrifugation at 30,000g. The supernatant was pipetted from the pellet and 20 mM 5′-(4-Fluorosulfonylbenzoyl)adenosine hydrochloride (FSBA; Sigma Aldrich) was added to the supernatant. The lysate was incubated with FSBA at 25 °C for 2 hours. Lysate was dialyzed in 2 L of assay buffer for 5 hours at room temperature followed by a second dialysis at 4 °C overnight in 2 L of assay buffer. The protein concentration in the lysate was then calculated with BCA assay Kit (∼9 mg/mL; Pierce).

Wilson et al.^29^ previously published the construction of an alignment and phylogenetic model using BAli-Phy^71^, which was used to resurrect ancestral protein sequences with PAML^72^. The robustness of these ancestral kinases have been previously tested by investigating activities of alternate sequences of the same nodes^29^. Ancestral and modern kinases were expressed and purified as previously reported^29^. The reaction was setup with 10 µM kinase, 20 mM MgCl_2_, 10 mM ATP, Phosphatase Inhibitor Cocktail #2 (Cal Biotech), and ∼1.3 mL of lysate (∼12 mg of protein). Reaction went for up to 4 hours at 25 °C. This was repeated three times for each kinase along with a background sample where no kinase was added. For the western dot blot time course of Src, 5 µL of sample was quenched with 15 µL of 8 M urea (Fisher Sci) to make the 4-fold dilutions. Other dilutions were made accordingly for 2-fold and 8-fold dilution samples. 1µL samples were loaded directly onto a nitrocellulose membrane (GenScript). The protocol was followed for the iBind Automated Western Systems (ThermoFisher) with Phosphotyrosine antibody (P-Tyr-1000 MultiMab(tm) Rabbit mAb mix; Cell Signaling Technology) as the primary antibody (1:2000 dilution) and ScanLater anti-rabbit antibody (Molecular Devices) as the secondary antibody (1:5000 dilution). The western dot blot was then imaged on a SpectraMax i3x Multi-Mode Microplate Reader with a ScanLater Western Blot cartridge (Molecular Devices).

### Mass Spectrometry of Phosphorylated Kinase-Inactive Lysate

For analysis of phosphorylated peptides, ∼1 mL of lysate (∼10 mg of protein) which has been reacted with the kinase previously was quenched with a 1:1 ratio of 8 M urea (Fisher). Followed by digestion with MS grade trypsin protease (Pierce) as instructed by the manufacturer. The reaction was then quenched with 10% trifluoroacetic acid (TFA, Sigma) bringing the final concentration to 0.5% TFA. Samples were desalted under vacuum using Sep Pak tC18 cartridges (Waters). Each cartridge was activated with 1 mL 80% acetonitrile (ACN)/0.1% TFA, then equilibrated with 3 × 1 mL of 0.1% TFA. Following sample loading, cartridges were washed with 4 × 1 mL of 0.1% TFA, and samples were eluted with 4 × 0.5 mL 50% ACN/0.25% formic acid (FA). 20 μg of each sample was kept for protein abundance measurements, and the remainder was used for phosphopeptide enrichment. Samples were dried by vacuum centrifugation. The digested peptides were enriched for phosphopeptides using ion metal affinity column (IMAC). FeCl_3_-NTA beads were prepared from Ni-NTA super flow slurry beads (QIAGEN) by first stripping the beads by incubating with 100 mM EDTA in a vacuum manifold three times. The beads were then washed with water before incubating with 15 mM FeCL_3_ (Sigma) for one minute three times. Excess FeCl_3_ was washed with water before rinsing the beads with 0.5% formic acid (FA). A slurry was prepared by adding water to the beads. 60 μL of slurry was added into a C18 NEST column that had been equilibrated with 150 μL of 80% ACN, 0.1% TFA. 100 μL of 50% ACN was added to the lyophilized lysate pellet to dissolve it. 100 μL of 100% ACN and 3 μL of 10% TFA was added after lysate was dissolved. 1 mg of peptide is added to Fe-NTA beads in the desalting tip and then incubated for 1-2 minutes followed by mixing and incubation for another 1-2 minutes. After incubating liquid is drained. Then 200 μL of 80% ACN and 0.1% TFA was used to wash the beads 3 times. 200 μL of 0.5% FA is used twice to wash the beads. the beads are incubated for 2-3 minutes with 200 μL of 500 mM phosphate buffer pH7 before eluting peptides to C18 column. This is repeated one more time to fully elute the peptides from the beads. The beads are incubated for 15 seconds with 200 μL 0.5% FA before the C18 column is used in a centrifuge to elute the phosphorylated peptides with 75 µL of 50% ACN and 0.1% TFA twice before the mass spectrometry run.

The LC/MS/MS was performed on all the samples prepared with phosphopeptide enrichment and a background sample with no enrichment. We used a 90-minute separation by nano reversed-phase HPLC gradient over a 75-um ID X 25-cm precolumn packed with Reprosil C18 1.9-μm (Waters). The sample was run on a Q-Exactive Plus mass spectrometer (ThermoFisher) and the top 20 ions were selected for MS2 sequencing. Resulting data was then searched with MaxQuant^73^ to against the human proteome to identify phosphorylated peptides. The mass spectrometry proteomics data have been deposited to the ProteomeXchange Consortium via the PRIDE partner repository^74^ with the dataset identifier PXD020299.

### Specificity Calculations

The results from MaxQuant^73^ were analyzed with an in-house script written in python. For each kinase, a set of substrate sequences was generated from the phosphorylated peptides found in at least one of the three trials. To generate the set of substrates, first the substrate peptides would be extended or shortened to 7 residues on each side of the phosphorylation site. If the sequence was too close to the beginning or end of a protein it would be rejected immediately. Next, if the sequence is already in the set of substrate sequences or was found in the control experiment, where no kinase was added, it would be rejected. Lastly, the localization probability must be greater than or equal to 70% and the MS intensity must be greater than 0. A background dataset was generated by applying the same rules to tyrosine containing peptides, from samples that were not enriched for phosphorylation. The background sequence logo was generated using WebLogo^75^.

Heatmaps were calculated by taking the log frequency of an amino acid occurring at a specific position minus the log of the frequency in the background. Each position and amino acid pair were tested using the logs odd ratio estimate used in pLogos and any significant value was marked with a black square in the heatmap.^*10*^ This allows for intensities of significant residues to be compared between data sets accurately. Residues below a 1.3-fold effect size were masked even if significant to focus only on residues which have the largest effect on specificity. Using the amino acid frequencies for each position the sequence entropy was calculated using the SciPy stats module^76^.

### Activity Assay

Initial rates were measured by a continuous colorimetric assay^77^. The reactions contained 20-100 nM of purified kinase along with 20 mM MgCl_2_ (Fisher), 525 µM β-Nicotinamide adenine dinucleotide (Sigma), 4 mM phosphoenolpyruvate (Sigma), 2.5 µL of PK/LDH (PK 600 U/mL – 1000 U/mL, LDH 900 U/mL-1400 U/mL; Sigma Aldrich), 0.3 mg/mL Bovine Serum Albumin (Fisher), and substrate peptide (GenScript). Reactions were initiated with 5 mM ATP, by pipetting the solution up and down, and then the absorbance was read at 340 nm for the course of the reaction (20 minutes) at 25 °C.

### Homology Model of Bound Peptide

An initial homology model was created from a crystal structure of Abl bound to an ATP-peptide conjugate (PDB: 2G2I). Using PyRosetta^78^, the initial structure was mutated to the sequence for either Src or Anc-S1. The bound peptide was then mutated to the sequence of Srctide. The backbone of the protein and peptide was set to be constrained before running the Fast Relax protocol using the ref2015 score function.

---

**Figure 1 — figure supplement 1:**
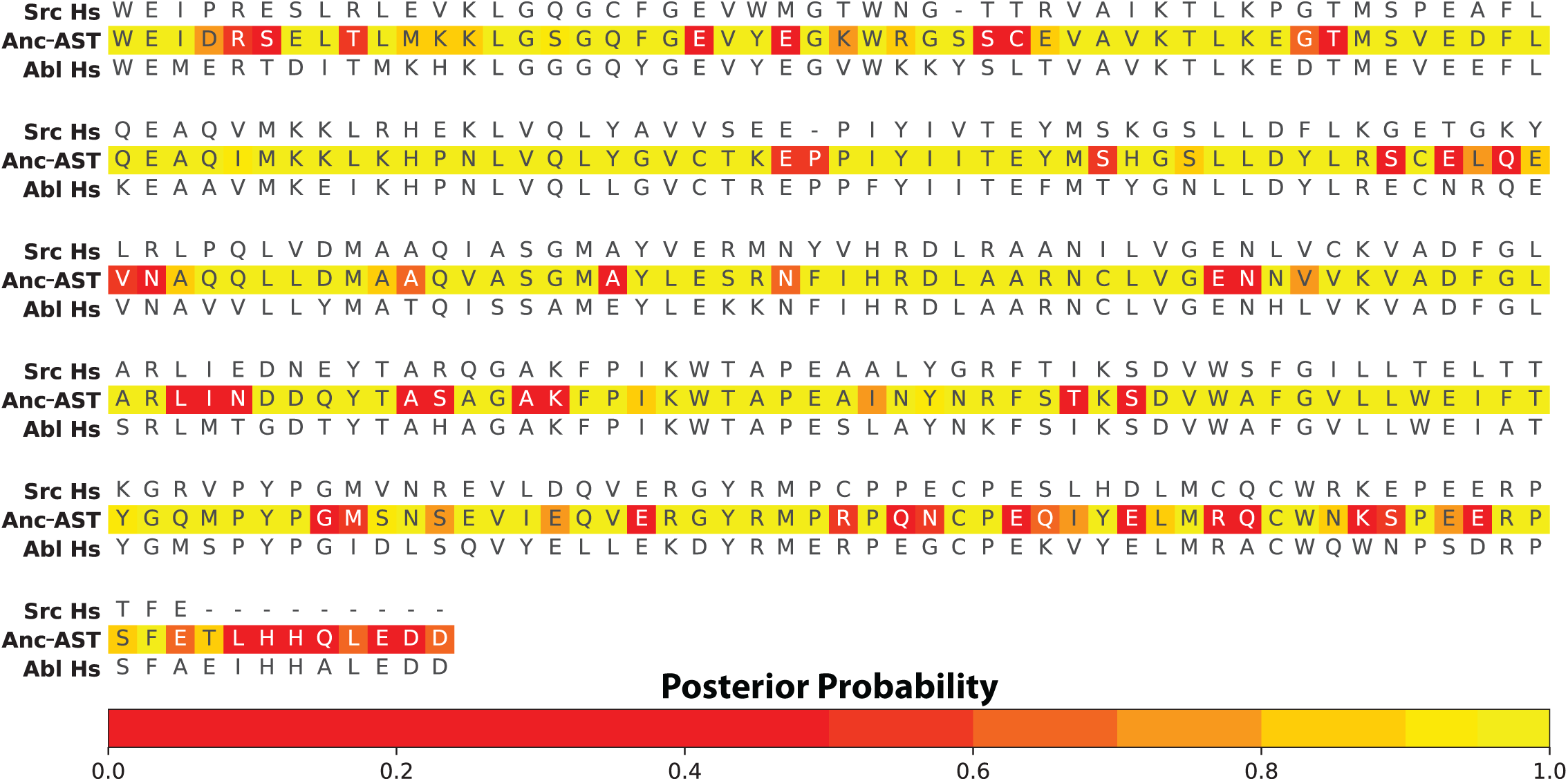
Posterior probabilities for Anc-AST reconstruction aligned with Src and Abl from *Homo sapiens*. Wilson *et al*. (2015) performed ancestral sequence reconstruction using PAML (Yang *et al*. 2007) and reported all other ancestral statistics and alignments. Here we report the Anc-AST sequence which has an average posterior probability of 0.86 across all positions. Anc-AST is aligned to the Src and Abl kinase domains and each position displays the given residue’s posterior probability from PAML.

**Figure 1 — figure supplement 2:**
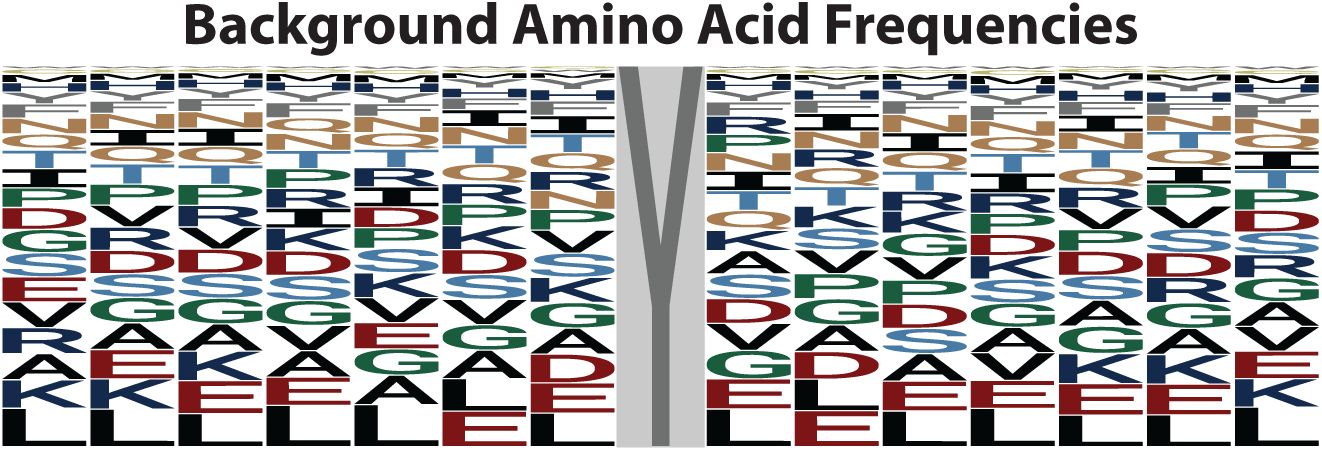
Frequencies of each amino acid from the unenriched background sample. Sequence logo generated by WebLogo (Crooks *et al*. 2004) displaying frequency of amino acids in peptides containing a phosphotyrosine from an unenriched sample, that also not treated with any target kinases.

*Figure 1 — figure supplement 3:* **BackgroundUnenrichedProteomics**.**csv** MaxQuant (Cox *et al*. 2008) results from a sample that was not enriched for phosphotyrosine. This data set was used to create the background amino acid frequencies.

*Figure 1— figure supplement 4:* **ControlPhosphoproteomicsData**.**csv** MaxQuant (Cox *et al*. 2008) results from phosphoproteomics experiment where no kinase was added.

**Figure 2 — figure supplement 1:**
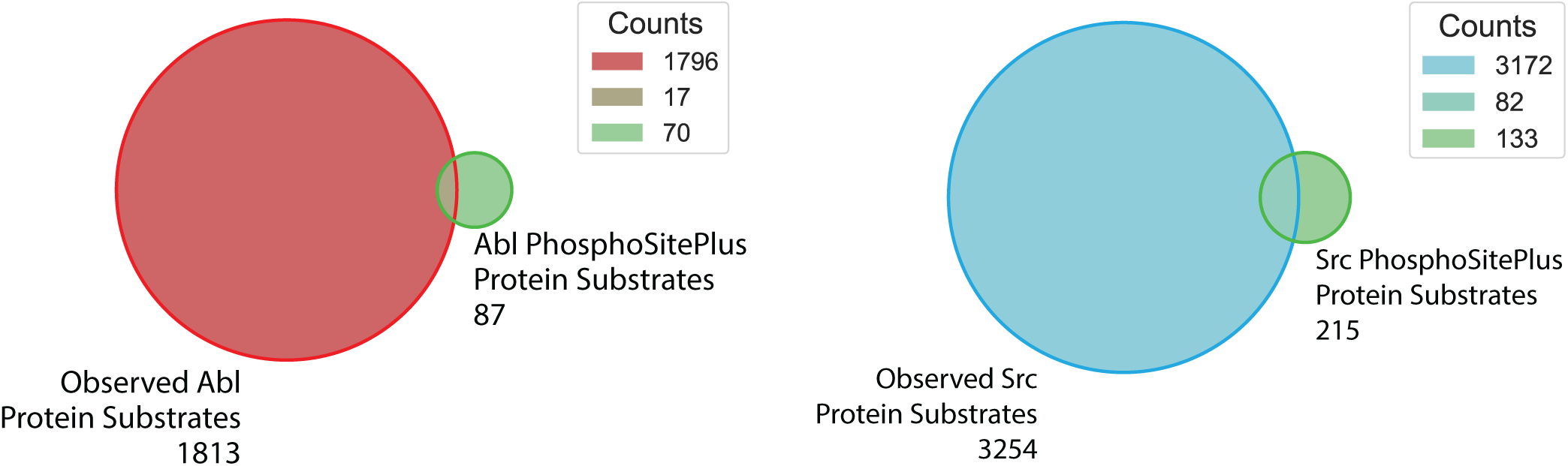
Natural protein substrates found in phosphoproteomics experiments. Venn diagrams of proteins that were observed in our kinase specificity experiments compared to the proteins listed in PhosphoSitePlus (Hornbeck *et al*. 2015) for the respective kinase.

**Figure 2 — figure supplement 2:**
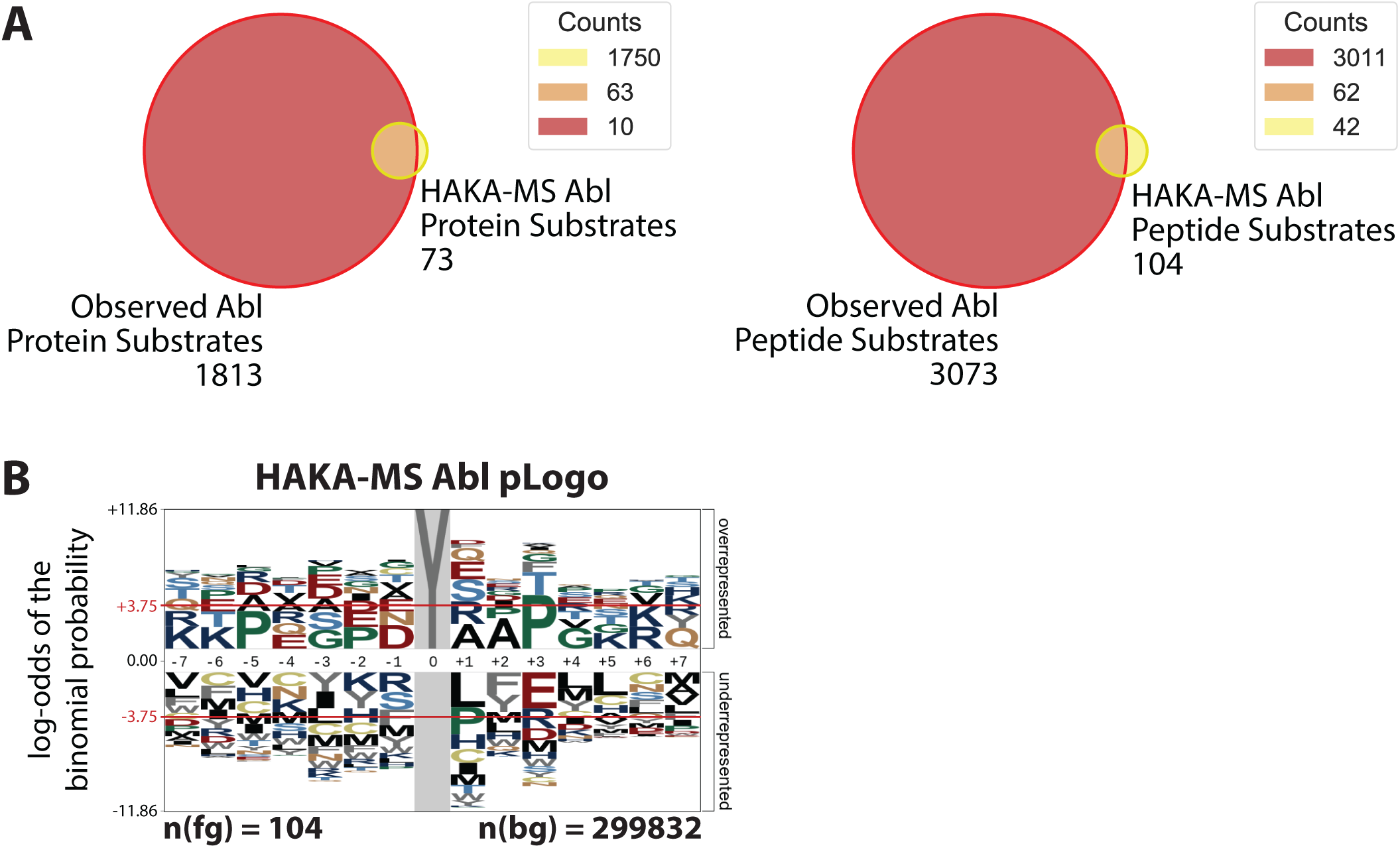
Comparison of our results with published data from the HAKA-MS experimental procedure (Muller *et al*. 2016). **(A)** Comparison of how many substrates (proteins and peptide fragments) were found in either our whole cell lysate assay or the HAKA-MS method. **(B)** pLogo showing the preference of Abl based on the peptides from the HAKA-MS method. The background used in their study was all tyrosines in the human proteome. Only P_+3_ was found to be statistically significant.

**Figure 3 — figure supplement 1:**
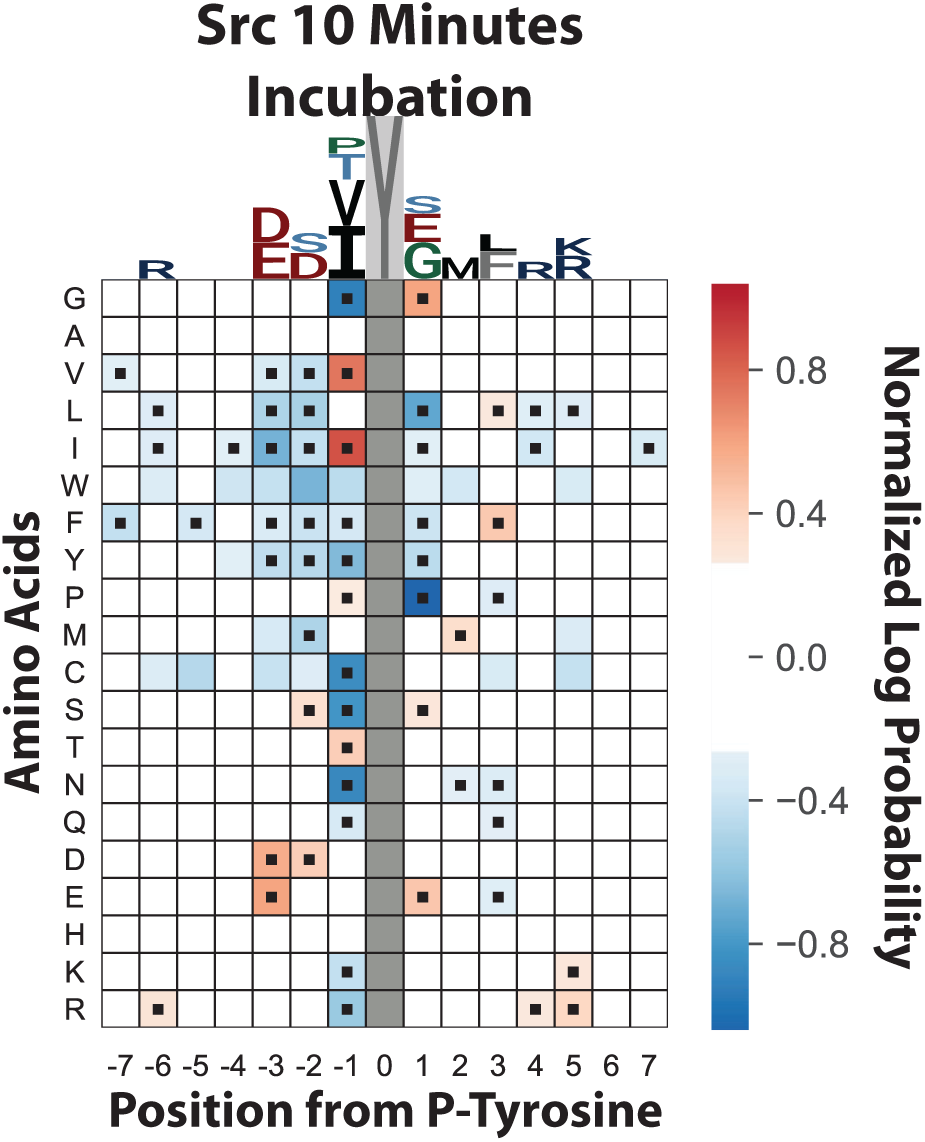
Substrate specificity is not affected by saturation phosphorylation. Heatmap displaying enrichment of amino acids at each position in a set of Src substrates after only 10 minutes of incubation. This short time point retains all major features of the heatmap from Src’s 4-hour time point in Figure 2C.

*Figure 3 — figure supplement 2:* **PhosphoproteomicsData.csv** MaxQuant results from phosphoproteomic experiments where kinase was added for the 4-hour incubation.

*Figure 3 — figure supplement 3:* **PhosphoproteomicsSrcShortTimePoint. csv** MaxQuant results rom the phosphoproteomic experiment where Src was added for only 10 minutes.

**Figure 5 — figure supplement 1:**
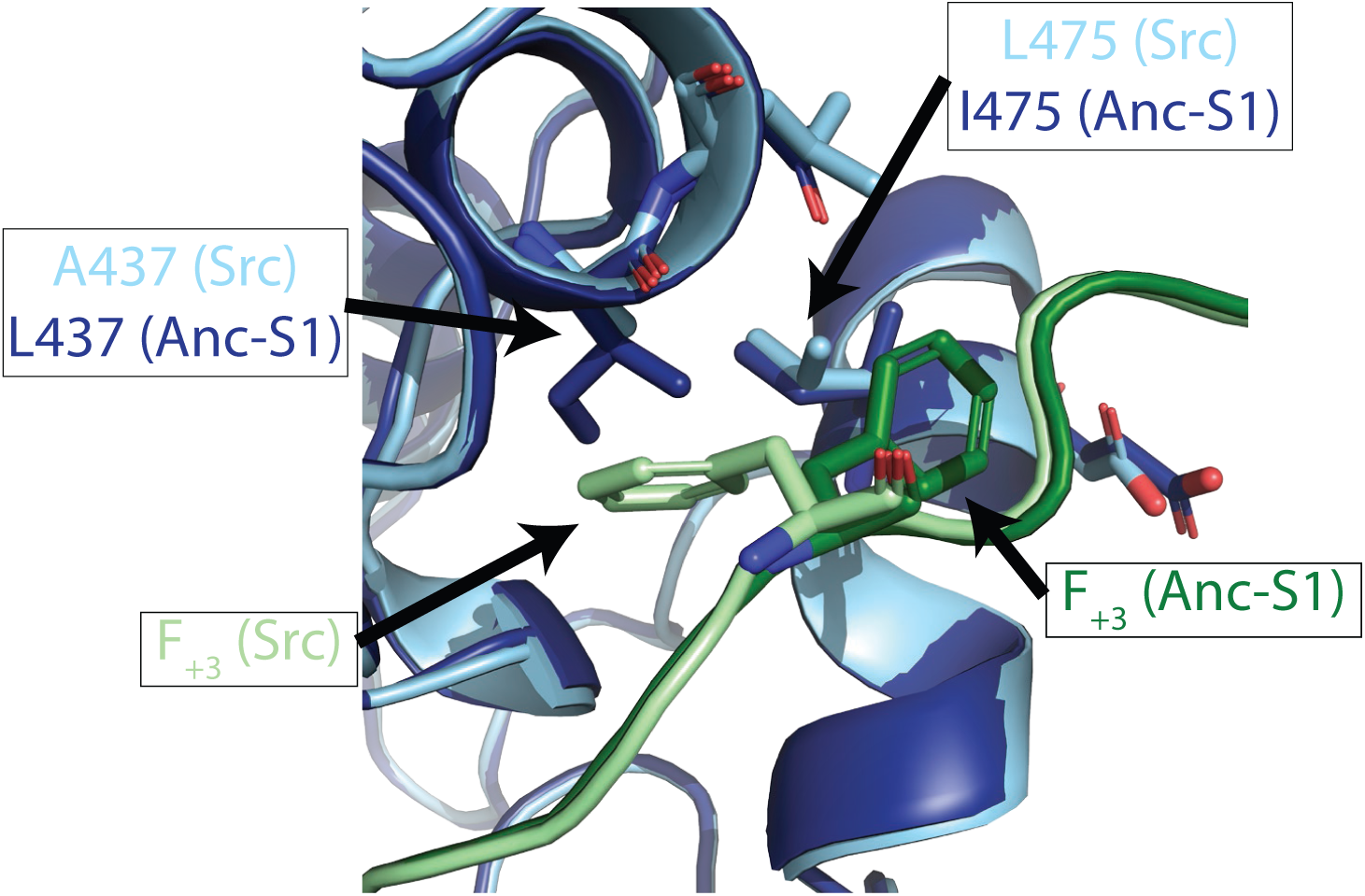
Homology modelling of protein/substrate complexes suggests amino acid differences between modern and ancestral proteins responsible for the differing substrate specificity in the +3 position. A crystal structure of Abl containing a peptide bound to the active site (PDB: 2G2I; Levinson *et al*. 2006) was used to build homology models of Src and Anc-S1 bound to Srctide. The bound peptide in PDB 2G2I already contained F_+3_, but the rest of the Srctide sequence was modelled in. A Fast Relax protocol was ran in Rosetta with full constraints on the backbone. The zoom-in indicates how the L475I and A437L substitutions in Anc-S1 could prohibit to bind F_+3_ in the same pocket as Src.

## References

1 Robinson, D. R., Wu, Y. M. & Lin, S. F. The protein tyrosine kinase family of the human genome. Oncogene 19, 5548–5557, doi:10.1038/sj.onc.1203957 (2000).

2 Parsons, S. J. & Parsons, J. T. Src family kinases, key regulators of signal transduction. Oncogene 23, 7906–7909, doi:10.1038/sj.onc.1208160 (2004).

3 Wang, J. Y. J. The Capable ABL: What Is Its Biological Function? Molecular and Cellular Biology 34, 1188–1197, doi:10.1128/mcb.01454-13 PMID - 24421390 (2014).

4 Ubersax, J. A. & Ferrell, J. E., Jr. Mechanisms of specificity in protein phosphorylation. Nat Rev Mol Cell Biol 8, 530–541, doi:10.1038/nrm2203 (2007).

5 Pinna, L. A. & Ruzzene, M. How do protein kinases recognize their substrates? Biochimica et Biophysica Acta (BBA) - Molecular Cell Research 1314, 191–225, doi:10.1016/s0167-4889(96)00083-3 (1996).

6 Mayer, B. J. & Baltimore, D. Mutagenic analysis of the roles of SH2 and SH3 domains in regulation of the Abl tyrosine kinase. Mol Cell Biol 14, 2883–2894, doi:10.1128/mcb.14.5.2883 (1994).

7 Mayer, B. J., Hirai, H. & Sakai, R. Evidence that SH2 domains promote processive phosphorylation by protein-tyrosine kinases. Current Biology 5, 296–305, doi:10.1016/s0960-9822(95)00060-1 (1995).

8 Yadav, S. S. & Miller, W. T. The evolutionarily conserved arrangement of domains in SRC family kinases is important for substrate recognition. Biochemistry 47, 10871–10880, doi:10.1021/bi800930e (2008).

9 Faux, M. C. & Scott, J. D. More on target with proteinphosphorylation: conferring specificity by location. Trends Biochem Sci 21, 312–315, doi:10.1016/s0968-0004(96)10040-2 PMID - 8772386 (1996).

10 Bhattacharyya, R. P., Remenyi, A., Yeh, B. J. & Lim, W. A. Domains, motifs, and scaffolds: the role of modular interactions in the evolution and wiring of cell signaling circuits. Annu Rev Biochem 75, 655–680, doi:10.1146/annurev.biochem.75.103004.142710 (2006).

11 Remenyi, A., Good, M. C. & Lim, W. A. Docking interactions in protein kinase and phosphatase networks. Curr Opin Struct Biol 16, 676–685, doi:10.1016/j.sbi.2006.10.008 (2006).

12 Hantschel, O. et al. Structural basis for the cytoskeletal association of Bcr-Abl/c-Abl. Mol Cell 19, 461–473, doi:10.1016/j.molcel.2005.06.030 (2005).

13 Songyang, Z. et al. Catalytic specificity of protein-tyrosine kinases is critical for selective signalling. Nature 373, 536–539, doi:10.1038/373536a0 (1995).

14 Till, J. H., Annan, R. S., Carr, S. A. & Miller, W. T. Use of synthetic peptide libraries and phosphopeptide-selective mass spectrometry to probe protein kinase substrate specificity. The Journal of Biological Chemistry 269, 7423–7428 (1994).

15 Scott, M. P. & Miller, W. T. A peptide model system for processive phosphorylation by Src family kinases. Biochemistry 39, 14531–14537, doi:10.1021/bi001850u (2000).

16 Schmitz, R., Baumann, G. & Gram, H. Catalytic specificity of phosphotyrosine kinases Blk, Lyn, c-Src and Syk as assessed by phage display. J Mol Biol 260, 664–677, doi:10.1006/jmbi.1996.0429 (1996).

17 Shah, N. H., Lobel, M., Weiss, A. & Kuriyan, J. Fine-tuning of substrate preferences of the Src-family kinase Lck revealed through a high-throughput specificity screen. Elife 7, e35190, doi:10.7554/eLife.35190 (2018).

18 Rychlewski, L., Kschischo, M., Dong, L., Schutkowski, M. & Reimer, U. Target specificity analysis of the Abl kinase using peptide microarray data. J Mol Biol 336, 307–311, doi:10.1016/j.jmb.2003.12.052 (2004).

19 Pawson, T. & Gish, G. D. SH2 and SH3 domains: from structure to function. Cell 71, 359–362, doi:10.1016/0092-8674(92)90504-6 (1992).

20 Pellicena, P., Stowell, K. R. & Miller, W. T. Enhanced phosphorylation of Src family kinase substrates containing SH2 domain binding sites. J Biological Chem 273, 15325–15328, doi:10.1074/jbc.273.25.15325 (1998).

21 Machiyama, H., Yamaguchi, T., Sawada, Y., Watanabe, T. M. & Fujita, H. SH3 domain of c-Src governs its dynamics at focal adhesions and the cell membrane. Febs J 282, 4034–4055, doi:10.1111/febs.13404 (2015).

22 Brehme, M. et al. Charting the molecular network of the drug target Bcr-Abl. Proc National Acad Sci 106, 7414–7419, doi:10.1073/pnas.0900653106 (2009).

23 Wu, J. J., Afar, D. E., Phan, H., Witte, O. N. & Lam, K. S. Recognition of multiple substrate motifs by the c-ABL protein tyrosine kinase. Comb Chem High Throughput Screen 5, 83–91, doi:10.2174/1386207023330516 (2002).

24 Hornbeck, P. V. et al. PhosphoSitePlus, 2014: mutations, PTMs and recalibrations. Nucleic Acids Res 43, D512–520, doi:10.1093/nar/gku1267 (2015).

25 Deng, Y. et al. Global analysis of human nonreceptor tyrosine kinase specificity using high-density peptide microarrays. J Proteome Res 13, 4339–4346, doi:10.1021/pr500503q (2014).

26 Harms, M. J. & Thornton, J. W. Evolutionary biochemistry: revealing the historical and physical causes of protein properties. Nat Rev Genet 14, 559–571, doi:10.1038/nrg3540 (2013).

27 Krishnan, N. M., Seligmann, H., Stewart, C. B., De Koning, A. P. & Pollock, D. D. Ancestral sequence reconstruction in primate mitochondrial DNA: compositional bias and effect on functional inference. Mol Biol Evol 21, 1871–1883, doi:10.1093/molbev/msh198 (2004).

28 Howard, C. J. et al. Ancestral resurrection reveals evolutionary mechanisms of kinase plasticity. Elife 3, doi:10.7554/eLife.04126 (2014).

29 Wilson, C. et al. Kinase dynamics. Using ancient protein kinases to unravel a modern cancer drug’s mechanism. Science 347, 882–886, doi:10.1126/science.aaa1823 (2015).

30 Benjamin, D. R. & Marc, A. S. Joint Bayesian Estimation of Alignment and Phylogeny. Systematic Biol 54, 401–418, doi:10.1080/10635150590947041 PMID - 16012107 (2005).

31 Williams, P. D., Pollock, D. D., Blackburne, B. P. & Goldstein, R. A. Assessing the accuracy of ancestral protein reconstruction methods. PLoS Comput Biol 2, e69, doi:10.1371/journal.pcbi.0020069 (2006).

32 Ferrari, S. et al. Aurora-A site specificity: a study with synthetic peptide substrates. Biochem J 390, 293–302, doi:10.1042/BJ20050343 (2005).

33 O’Shea, J. P. et al. pLogo: a probabilistic approach to visualizing sequence motifs. Nat Methods 10, 1211–1212, doi:10.1038/nmeth.2646 (2013).

34 Knight, J. D. et al. A novel whole-cell lysate kinase assay identifies substrates of the p38 MAPK in differentiating myoblasts. Skelet Muscle 2, 5, doi:10.1186/2044-5040-2-5 (2012).

35 Muller, A. C. et al. Identifying Kinase Substrates via a Heavy ATP Kinase Assay and Quantitative Mass Spectrometry. Sci Rep 6, 28107, doi:10.1038/srep28107 (2016).

36 Schwartz, D. & Gygi, S. P. An iterative statistical approach to the identification of protein phosphorylation motifs from large-scale data sets. Nat Biotechnol 23, 1391–1398, doi:10.1038/nbt1146 (2005).

37 Fuchs, J. E. et al. Cleavage entropy as quantitative measure of protease specificity. PLoS Comput Biol 9, e1003007, doi:10.1371/journal.pcbi.1003007 (2013).

38 Shah, N. H. et al. An electrostatic selection mechanism controls sequential kinase signaling downstream of the T cell receptor. Elife 5, e20105, doi:10.7554/eLife.20105 (2016).

39 Foda, Z. H., Shan, Y., Kim, E. T., Shaw, D. E. & Seeliger, M. A. A dynamically coupled allosteric network underlies binding cooperativity in Src kinase. Nat Commun 6, 5939, doi:10.1038/ncomms6939 (2015).

40 Colicelli, J. ABL tyrosine kinases: evolution of function, regulation, and specificity. Sci Signal 3, re6, doi:10.1126/scisignal.3139re6 (2010).

41 Till, J. H., Chan, P. M. & Miller, W. T. Engineering the substrate specificity of the Abl tyrosine kinase. J Biological Chem 274, 4995–5003, doi:10.1074/jbc.274.8.4995 (1999).

42 Moses, A. M. & Landry, C. R. Moving from transcriptional to phospho-evolution: generalizing regulatory evolution? Trends Genet 26, 462–467, doi:10.1016/j.tig.2010.08.002 (2010).

43 Manning, G., Whyte, D. B., Martinez, R., Hunter, T. & Sudarsanam, S. The protein kinase complement of the human genome. Science 298, 1912–1934, doi:10.1126/science.1075762 (2002).

44 Copley, S. D. Evolution of new enzymes by gene duplication and divergence. Febs J 287, 1262–1283, doi:10.1111/febs.15299 (2020).

45 Conant, G. C. & Wolfe, K. H. Turning a hobby into a job: How duplicated genes find new functions. Nature Reviews Genetics 9, 938–950, doi:10.1038/nrg2482 PMID - 19015656 (2008).

46 Innan, H. & Kondrashov, F. The evolution of gene duplications: classifying and distinguishing between models. Nat Rev Genetics 11, 97–108, doi:10.1038/nrg2689 PMID - 20051986 (2010).

47 Muller, H. J. Bar Duplication. Science 83, 528–530, doi:10.1126/science.83.2161.528-a (1936).

48 Ohno, S. Evolution by Gene Duplication. doi:10.1007/978-3-642-86659-3 (1970).

49 Force, A. et al. Preservation of duplicate genes by complementary, degenerative mutations. Genetics 151, 1531–1545 (1999).

50 Hughes, A. L. The evolution of functionally novel proteins after gene duplication. Proc Biol Sci 256, 119–124, doi:10.1098/rspb.1994.0058 (1994).

51 Lynch, M. & Conery, J. S. The evolutionary fate and consequences of duplicate genes. Science 290, 1151–1155, doi:10.1126/science.290.5494.1151 (2000).

52 Walsh, J. B. How often do duplicated genes evolve new functions? Genetics 139, 421–428 (1995).

53 Wheeler, L. C., Anderson, J. A., Morrison, A. J., Wong, C. E. & Harms, M. J. Conservation of Specificity in Two Low-Specificity Proteins. Biochemistry 57, 684–695, doi:10.1021/acs.biochem.7b01086 (2018).

54 Siddiq, M. A., Hochberg, G. K. & Thornton, J. W. Evolution of protein specificity: insights from ancestral protein reconstruction. Curr Opin Struct Biol 47, 113–122, doi:10.1016/j.sbi.2017.07.003 (2017).

55 Boucher, J. I., Jacobowitz, J. R., Beckett, B. C., Classen, S. & Theobald, D. L. An atomic-resolution view of neofunctionalization in the evolution of apicomplexan lactate dehydrogenases. Elife 3, e02304, doi:10.7554/eLife.02304 (2014).

56 McKeown, A. N. et al. Evolution of DNA specificity in a transcription factor family produced a new gene regulatory module. Cell 159, 58–68, doi:10.1016/j.cell.2014.09.003 (2014).

57 Aakre, C. D. et al. Evolving new protein-protein interaction specificity through promiscuous intermediates. Cell 163, 594–606, doi:10.1016/j.cell.2015.09.055 PMID - 26478181 (2015).

58 Pellicena, P. & Miller, W. T. Processive phosphorylation of p130Cas by Src depends on SH3-polyproline interactions. J Biological Chem 276, 28190–28196, doi:10.1074/jbc.M100055200 (2001).

59 Filippakopoulos, P. et al. Structural coupling of SH2-kinase domains links Fes and Abl substrate recognition and kinase activation. Cell 134, 793–803, doi:10.1016/j.cell.2008.07.047 (2008).

60 Lorenz, S., Deng, P., Hantschel, O., Superti-Furga, G. & Kuriyan, J. Crystal structure of an SH2-kinase construct of c-Abl and effect of the SH2 domain on kinase activity. Biochem J 468, 283–291, doi:10.1042/BJ20141492 (2015).

61 Songyang, Z. et al. Specific motifs recognized by the SH2 domains of Csk, 3BP2, fps/fes, GRB-2, HCP, SHC, Syk, and Vav. Mol Cell Biol 14, 2777–2785, doi:10.1128/mcb.14.4.2777 (1994).

62 Lim, W. A., Richards, F. M. & Fox, R. O. Structural determinants of peptide-binding orientation and of sequence specificity in SH3 domains. Nature 372, 375–379, doi:10.1038/372375a0 (1994).

63 Bradshaw, J. M., Mitaxov, V. & Waksman, G. Mutational investigation of the specificity determining region of the src SH2 domain 1 1Edited by J. A. Wells. Journal of Molecular Biology 299, 523–537, doi:10.1006/jmbi.2000.3765 PMID - 10860756 (2000).

64 Marengere, L. E. et al. SH2 domain specificity and activity modified by a single residue. Nature 369, 502–505, doi:10.1038/369502a0 (1994).

65 Waksman, G. & Kuriyan, J. Structure and specificity of the SH2 domain. Cell 116, S45-48, 43 p following S48, doi:10.1016/s0092-8674(04)00043-1 (2004).

66 Ren, R., Ye, Z. S. & Baltimore, D. Abl protein-tyrosine kinase selects the Crk adapter as a substrate using SH3-binding sites. Gene Dev 8, 783–795, doi:10.1101/gad.8.7.783 PMID - 7926767 (1994).

67 Moran, M. F. et al. Src homology region 2 domains direct protein-protein interactions in signal transduction. Proc National Acad Sci 87, 8622–8626, doi:10.1073/pnas.87.21.8622 (1990).

68 Boggon, T. J. & Eck, M. J. Structure and regulation of Src family kinases. Oncogene 23, 7918–7927, doi:10.1038/sj.onc.1208081 (2004).

69 Shah, N. H., Amacher, J. F., Nocka, L. M. & Kuriyan, J. The Src module: an ancient scaffold in the evolution of cytoplasmic tyrosine kinases. Crit Rev Biochem Mol Biol 53, 535–563, doi:10.1080/10409238.2018.1495173 (2018).

70 Miller, W. T. Determinants of substrate recognition in nonreceptor tyrosine kinases. Acc Chem Res 36, 393–400, doi:10.1021/ar020116v (2003).

71 Suchard, M. A. & Redelings, B. D. BAli-Phy: simultaneous Bayesian inference of alignment and phylogeny. Bioinformatics 22, 2047–2048, doi:10.1093/bioinformatics/btl175 (2006).

72 Yang, Z. PAML 4: phylogenetic analysis by maximum likelihood. Mol Biol Evol 24, 1586–1591, doi:10.1093/molbev/msm088 (2007).

73 Cox, J. & Mann, M. MaxQuant enables high peptide identification rates, individualized p.p.b.-range mass accuracies and proteome-wide protein quantification. Nat Biotechnol 26, 1367–1372, doi:10.1038/nbt.1511 (2008).

74 Perez-Riverol, Y. et al. The PRIDE database and related tools and resources in 2019: improving support for quantification data. Nucleic acids research 47, D442–D450, doi:10.1093/nar/gky1106 PMID - 30395289 (2018).

75 Crooks, G. E., Hon, G., Chandonia, J.-M. & Brenner, S. E. WebLogo: A Sequence Logo Generator. Genome Research 14, 1188–1190, doi:10.1101/gr.849004 PMID - 15173120 (2004).

76 Virtanen, P. et al. SciPy 1.0--Fundamental Algorithms for Scientific Computing in Python. Arxiv 17, 261–272, doi:10.1038/s41592-019-0686-2 PMID - 32015543 (2019).

77 Barker, S. C. et al. Characterization of pp60c-src tyrosine kinase activities using a continuous assay: autoactivation of the enzyme is an intermolecular autophosphorylation process. Biochemistry 34, 14843–14851, doi:10.1021/bi00045a027 (1995).

78 Chaudhury, S., Lyskov, S. & Gray, J. J. PyRosetta: a script-based interface for implementing molecular modeling algorithms using Rosetta. Bioinformatics 26, 689–691, doi:10.1093/bioinformatics/btq007 PMID - 20061306 (2010).

